# iSparse kmeans: a two-step clustering approach for big dynamic functional network connectivity data

**DOI:** 10.1101/2022.03.13.484193

**Authors:** Mohammad S. E. Sendi, David H Salat, Robyn L Miller, Vince D Calhoun

## Abstract

**Background:** Dynamic functional network connectivity (dFNC) estimated from resting-state functional magnetic imaging (rs-fMRI) studies the temporally varying of functional integration between brain networks. In a typical dFNC pipeline, a clustering stage to summarize the connectivity patterns that are transiently but reliably realized over the course of a scanning session. However, identifying the right number of clusters through a conventional clustering criterion computed by running the algorithm repeatedly, over a large range of cluster numbers is time-consuming and requires substantial computational power even for typical dFNC datasets, and the computational demands become prohibitive as datasets become larger and scans longer. Here we developed a new dFNC pipeline, called iterative sparse kmeans or iSparse kmeans, to analyze large dFNC data without having access to huge computational power.

**Method:** In iSparse kmeans, we implement two-step clustering. In the first step, we randomly use a sub-sample dFNC data and identify several sets of states at different model orders. In the second step, we aggregate all dFNC states estimated from all iterations in the first step and use this to identify the optimum number of clusters using the elbow criteria. Additionally, we use this new reduced dataset and estimate a final set of states by performing a second kmeans clustering on the aggregated dFNC states from the first k-means clustering. To validate the reproducibility of iSparse kmeans, we analyzed four dFNC datasets from the human connectome project (HCP).

**Results:** We found that both conventional kmeans and iSparse kmeans generate similar brain dFNC states while iSparse kmeans is 27 times faster than the traditional method in finding the optimum number of clusters. We show that the results are replicated across four different datasets from HCP.

**Conclusion:** We developed a new analytic pipeline which facilitates analysis of large dFNC datasets without having access to a huge computational power source. We validated the reproducibility of the result across multiple datasets.

## 1. Introduction

In recent decades, blood-oxygenation-level-dependent (BOLD) functional magnetic resonance imaging (fMRI) has provided unique information about brain changes associated with various brain disorders (Heeger and Ress, 2002; Poldrack, 2008; Carbó-Carreté et al., 2020). fMRI is a non-invasive imaging technique that identifies localized, time-varying alterations in brain metabolism, such as blood flow and deoxygenated hemoglobin levels (Herberholz et al., 2011). These metabolic changes can be induced by a cognitive task (i.e., task-based fMRI) (Cook et al., 2020) or via unregulated brain fluctuations during rest (i.e., resting-state fMRI). Functional connectivity (FC) or its network analog functional network connectivity (FNC) studies the temporal dependence (typically assessed with correlation) between the BOLD fMRI signal from different brain regions (van den Heuvel and Hulshoff Pol, 2010). The FNC approach uses temporal dependence to infer how various brain networks communicate and may play a significant role in understanding how large-scale neuronal communication in the human brain relates to human behavior (Kalinosky et al., 2019; Cook et al., 2020) and how neurodegenerative diseases alter this relationship (Wang et al., 2019; Yan et al., 2019; Hummer et al., 2020; Quevenco et al., 2020; Vega et al., 2020).

Most previous studies assume FNC is static over time and ignore (average out) brain dynamics (Ioannides, 2007). Indeed, functional connectivity is highly dynamic, even during the resting state (Sendi et al., 2021c). In recent years, a new line of research called dynamic functional network connectivity (dFNC) have moved beyond studying the strength of connectivity among brain regions and studied the temporal properties of the FNC (Allen et al., 2014). dFNC has shown promise as a biomarker for schizophrenia (Sendi et al., 2021a, 2021b), Alzheimer’s disease (Sendi et al., 2021c), major depressive disorder (Sendi et al., 2021e), and autism spectrum disorder (Harlalka et al., 2019). It has been shown that dFNC improves classification between disordered and healthy conditions (Rashid et al., 2015; Saha et al., 2021) and provides more information about the pathology of neurological and neuropsychiatric disorders than its static counterpart (Menon and Krishnamurthy, 2019).

Fig.1 shows the analytic pipeline that is used for analyzing dFNC information(Rashid et al., 2015; Sendi et al., 2021c, 2021b, 2021a, 2021e). This pipeline contains four main steps. In the first step, we estimate the intrinsic components for the desired brain regions. Second, we calculate the dFNC using a sliding window. In the third step, we concatenate all dFNCs of all subjects and go through an optimization process to find the clustering order based on the elbow criterion. In the fourth step, we estimate the final dFNC for the whole group and state vector for each individual and calculate the dFNC features for statistical analysis.

**Fig.1.**
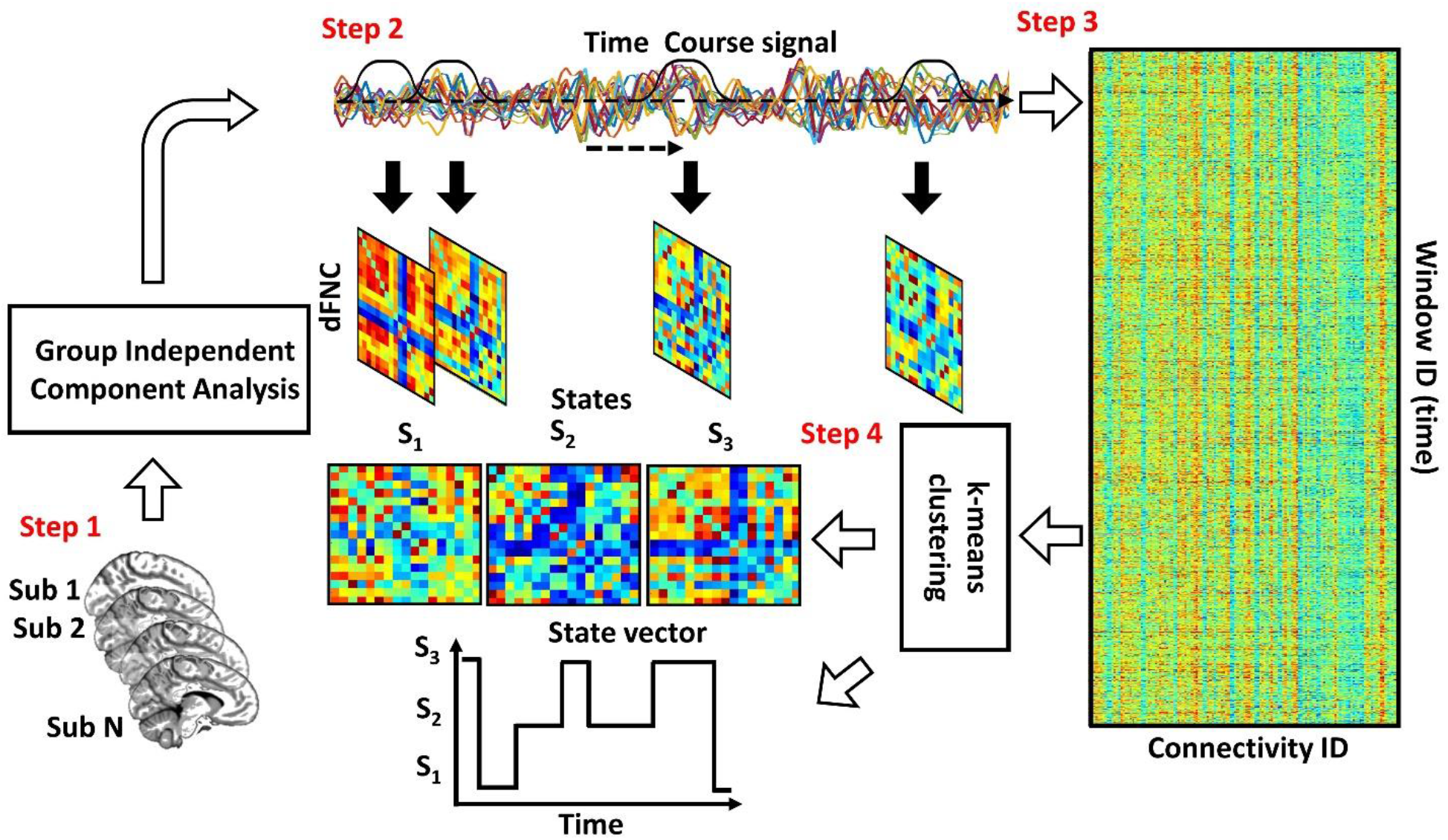
The conventional dFNC pipeline. In Step1, we estimate the independent components using group independent component analysis. In Step2, we estimate the dFNC using sliding window. In Step3, we concatenate all dFNCs across all participants. Then, based on elbow criteria, we estimate the cluster order. In step4, we use a standard kmeans clustering approach and calculate the dFNC state for group and state vector for everyone.

Even though any clustering approach can be used for clustering dFNC information, mainly k-means clustering has been used due to its simplicity in implementation, ability to scale to a large dataset (Fränti and Sieranoja, 2019). Additionally, it has been shown that kmeans clustering is faster than the other methods such as spectral clustering, density-based spatial clustering of applications with noise or DBSCAN, and mean-shift clustering (McInnes and Healy, 2017). But it is still slow and needs substantial computational power when we work on a sizeable dFNC dataset. On the other hand, recently, the availability of extremely large neuroimaging datasets has made the computational burden of clustering of dFNC measurements a significant practical challenge. For example, the UK Biobank dataset released neuroimaging data from more than 40,000 participants (Alfaro-Almagro et al., 2021) and has targeted acquiring data from 100,000 individuals (Alfaro-Almagro et al., 2018). Also, it has been discussed that many neuroimaging analytic pipelines are not scalable for massive data sets, including possibly tens, if not hundreds of thousands of participants (Van Horn and Toga, 2014). Therefore, developing a framework that can analyze a large dFNC dataset within a reasonable timeframe in a typical cluster computing environment is needed.

We introduce a new iterative clustering algorithm, iterative sparse k-means (iSparse k-means), that efficiently scales to millions of high-dimensional observations, making it a valuable addition to the pipeline for large scale dFNC analyses. We evaluated the reproducibility of the results with both standard and proposed dFNC pipelines across four rs-fMRI sessions of HCP young adults. Additionally, we compared the time is needed to find the optimal cluster number with iSparse kmeans versus standard kmeans, and showed our approach is faster than the standard method in finding the cluster order.

## 2. Material and methods

Our analytic pipeline includes rs-fMRI preprocessing, extracting independent components, calculating dFNC, estimating the cluster order and dFNC states using the proposed clustering method. The following subsection describes each step in more detail.

### 2.1. Preprocessing and independent components extraction

We used the statistical parametric mapping (SPM12, https://www.fil.ion.ucl.ac.uk/spm/) running in MATLAB2019 to preprocess the fMRI data. The first five dummy scan were removed before preprocessing. Rigid body motion correction was used to account for participant’s head movement. Then, we used spatial normalization by echo-planar imaging (EPI) template into the standard Montreal Neurological Institute (MNI) space. Finally, a Gaussian kernel was used to smooth the fMRI images using a full width at half maximum (FWHM) of 6mm. Next, we adapted Neuromark pipeline to extract intrinsic connectivity networks (ICNs) for each subject (Du et al., 2020). Using this pipeline, we estimated 53 ICNs for each subject and categorized them into seven network domains, including subcortical network (SCN), auditory network (ADN), sensorimotor network (SMN), visual network (VSN), cognitive control network (CCN), the default-mode network (DMN), and cerebellar network (CBN) as shown in Fig. 2. The details of the extracted ICNs are provided in (Sendi et al., 2021d).

**Fig.2.**
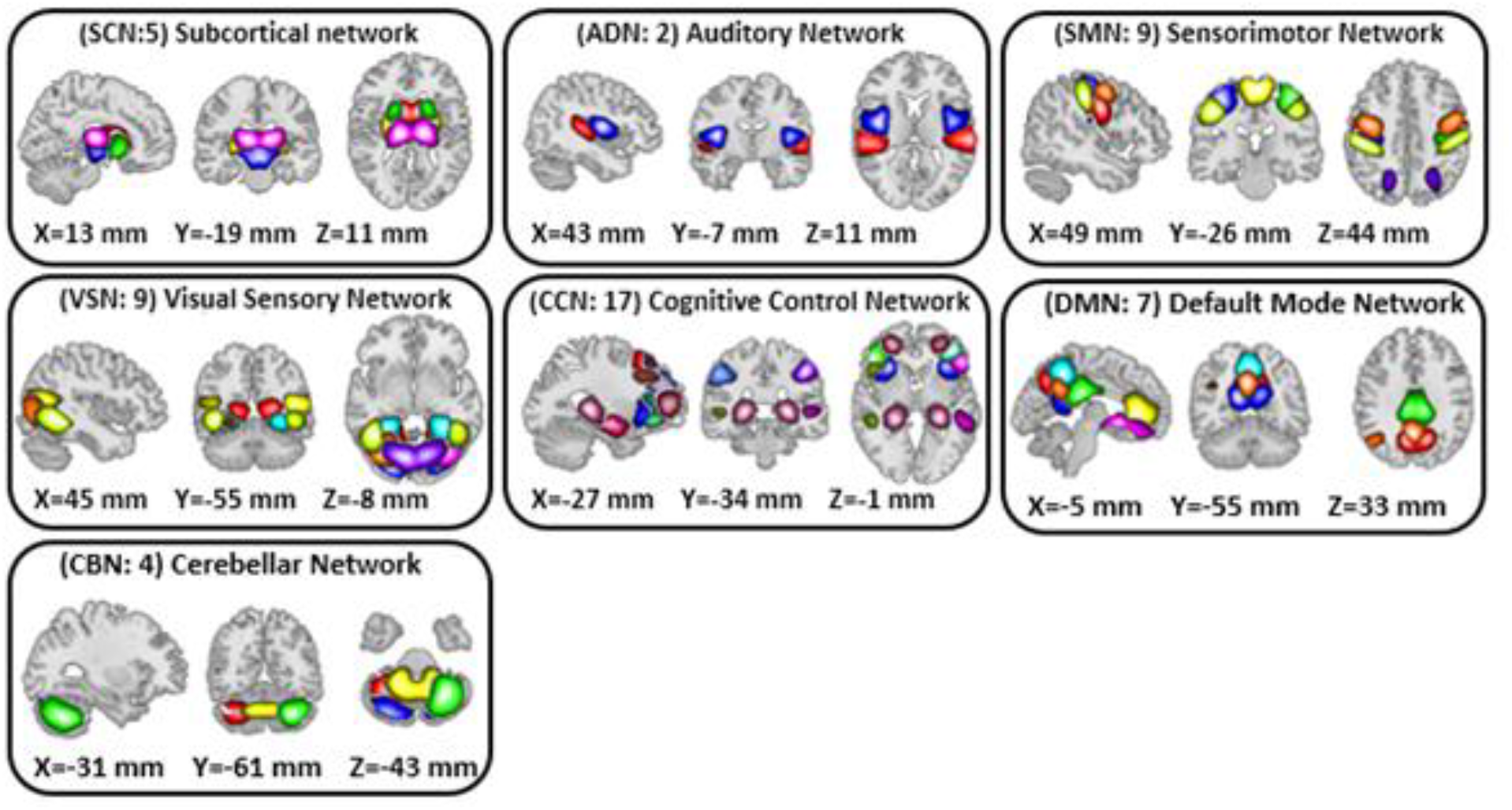
Extracted independent components. 53 independent components estimated by NeuroMark pipeline. We put them in seven domains including subcortical network (SCN), auditory network (AND), sensorimotor network (SMN), visual sensory network (VSN), cognitive control network (CCN), default mode network (DMN). and cerebellar network (CBN).

### 2.2. Dynamic functional network connectivity estimation

We used a tapered sliding window and estimated the functional connectivity within each window using the Pearson correlation as shown in Eq.1.

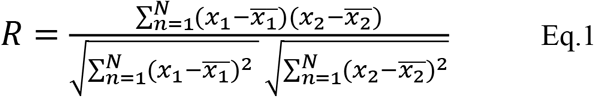

where *x*_1_ and *x*_2_ are time-course signals and 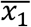 and 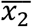 are the mean of *x*_1_ and *x*_2_, respectively. It takes values in the interval [− 1, 1] and measures the strength of the linear relationship between *x*_1_ and *x*_2_.

With 53 ICN, the size of each dFNC is 53 × 53, which equals 1378 distinct connectivity features. Next, we concatenated dFNC estimates of each window for each subject to form a matrix, called dFNC tensor hereafter, with the size of *T*× *F*, where T denotes the number of windows and F donates the number of connectivity features (Fig.3).

**Fig.3.**
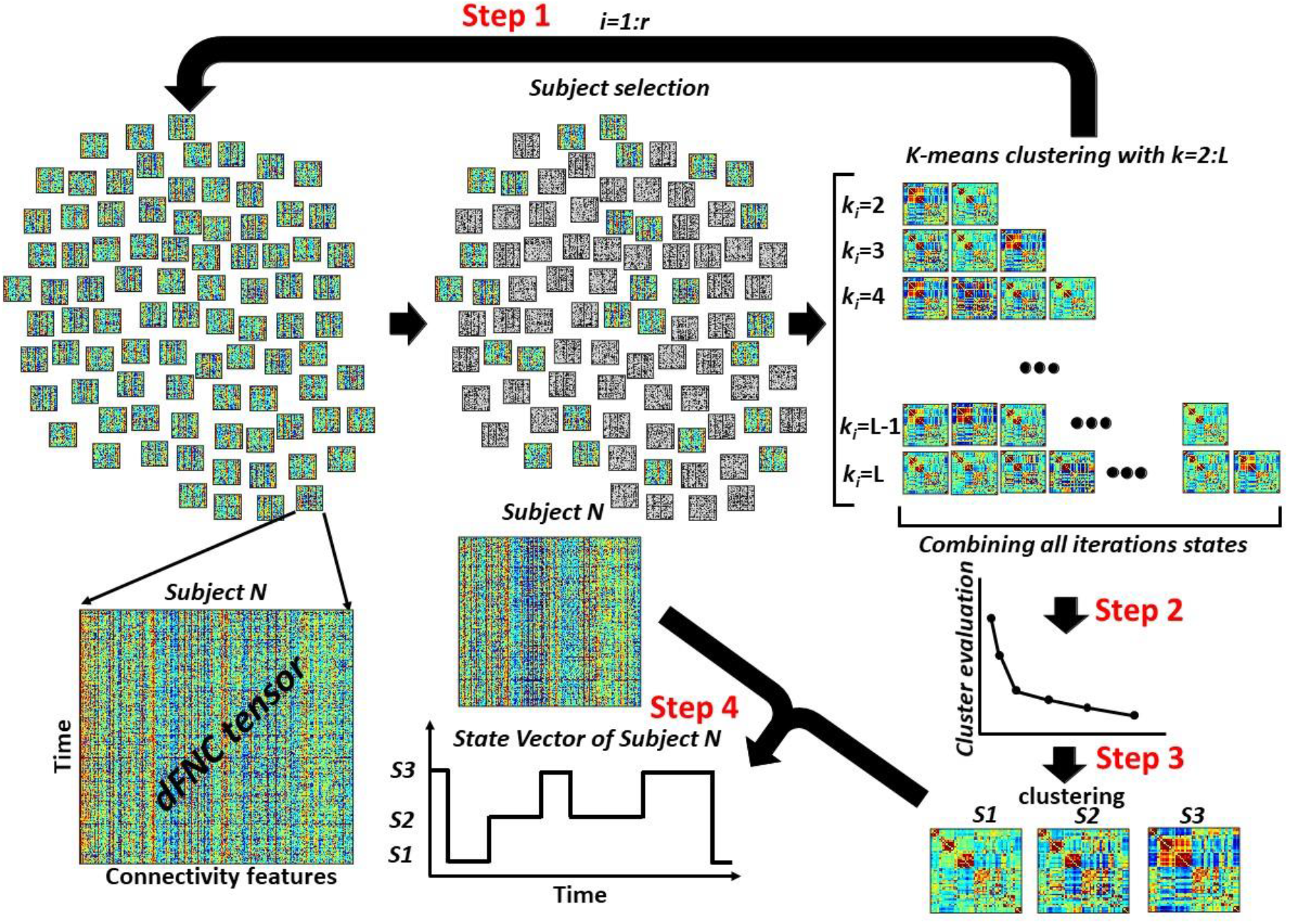
The overview of iSparse Kmeans clustering approach for dFNC state estimation. In Step 1, we select a subsample of dFNC tensor and then used kmeans clustering with k values from 2 to L and put them into 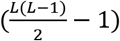. With *r* iteration, we would have 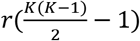 clusters centroids in 2, concatenated all cluster centroids and we use elbow criteria ttootfainl.dInthSetebpest k values, called K_opt_ hereafter. In Step3, using another kmeans clustering approach, we estimated the final dFNC states. In Step 4, we used this final states and found the state vector for each subject.

### 2.3. iSparse kmeans clustering

Fig. 3 shows the proposed iSparse Kmeans clustering method for estimating dFNC states. This method includes a few steps. **Step1:** We sub-sample subjects dFNC tensors (*m* subjects from *n* subjects per iteration). Then, we run a standard kmeans clustering on the subsampled data with different values of *k* = 2,3,.., *L*. The k-means algorithm divides *m* × *T* samples *X* of each iteration into *k* disjoint clusters *C*_1_, *C*_2_,…, *C*_*k*_. The cluster centroids *μ*_*i*_ of *C*_*i*_ minimize the within-cluster sum-of-squares criterion as shown in Eq.2.

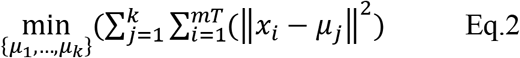

We exhaust all subjects by repeating this process *r* times over disjoint sets of *m* subjects, where *r* is equal to 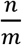. In each iteration, we save all cluster centroids for all values of *k* ∈ [2, *L*] Therefore, we would have 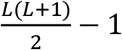 representative cluster centroids in each iteration. By repeating this process *r* times, we would have 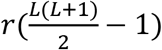 cluster centroids, a reduction of the data from the whole dFNC. **Step 2:** We concatenate all centroids estimated from all *r* iterations.Next, we use the elbow criteria to find the optimum number of clusters using all 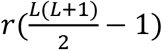 observations. **Step 3:** After finding the optimum number of clusters, called *K*_*opt*_ hereafter, we use another standard k-means clustering to put all 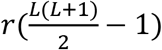 states into K_opt_ cluster, called final states. **Step 4:** Using the final *K*_*opt*_ states, we assign the dFNC of each subject to one of the estimated states and extract the state vector of each participant.

### 2.4. dFNC temporal features estimation

We estimated the occupancy rate (OCR) and the number of transitions between states as the representative dFNC temporal features from the state vector. The OCR represents the proportional amount of time each individual spends in a given state for all HCP datasets through both standard and isparse kmeans methods.

### 2.5. Clustering quality assessment

To assess the clustering quality for each dFNC data, we calculated the distance between the dFNC data and its associated cluster centroid. Then we calculated the distance between each dFNC sample with the other cluster centroids and then summed them up. Then, we calculated the ratio of the latter to the former one for each dFNC instance, called distance ratio here. Finally, we averaged all distance ratios out for each participant.

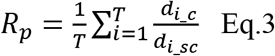

*d*_*ic*_ is the distance between each sample to the cluster centroid of the state the sample belongs. Also, *d*_*ij*_ is the distance between each sample to other cluster centroids. *r*_*i*_ is the distance ratio for each sample and *R*_*p*_ is the averaged distance ratio for each participant. It is worth mentioning that a higher ratio means better quality in clustering.

### 2.6. Dataset

To test the proposed method, we used the rs-fMRI and demographic information collected from the 833 young healthy adults (average age: 28.65; range: 22-37 years; female/male: 443/390) from the Human Connectome Project (HCP) (Glasser et al., 2016). This dataset is available on the HCP website (https://www.humanconnectome.org). The institutional review board from both Washington University and University of Minnesota approved the study. The rs-fMRI data were collected on a Siemens Skyra 3T with a 32-channel RF receiver head coil. High resolution T2*- weighted functional images were acquired using a gradient-echo EPI sequence with TE = 33.1 ms, TR= 0.72 s, flip angle = 52°, slice thickness = 2 mm, 72 s slices and 2 mm isotropic voxel, the field of view: 208×180 mm (RO×PE), and duration: 14:33 (min: sec). For each participant, four separated rs-fMRI sessions (two sessions per day) were acquired that are called HCP1 (session1, day1), HCP2 (session2, day1), HCP3 (session 1, day2), and HCP4 (session2, day12) hereafter. We used all four sessions to evaluate the reproducibility of the result using the proposed dFNC states estimation method. The dFNC size of HCP1, HCP2, HCP3, and HCP4 is 848827×1378 (8542 MB), 732207×1378 (7403 MB), 747201×1378 (7555 MB), and 769692×1378 (7742 MB), respectively.

## 3. Results

### 3.1. Standard kmeans and iSparse kmeans clustering produce similar brain states

The first question we were interested in answering is whether both standard kmeans and iSparse kmeans would generate similar dFNC states or not. To test this, we clustered the dFNC data with different *L* values in iSparse kmeans (as shown in Fig.3). In the iSparse kmeans, we used 3% of the entire dataset in each iteration. Using elbow criteria, we found that the optimal number of clusters is 2 through both standard and proposed kmeans clustering approach. Then, to evaluate the similarity of dFNC states estimated by iSparse kmeans (with different L) with the states estimated by conventional kmeans, we used the correlation across the matched states as a similarity metric. The similarity between matched states with varying values of L is shown in Fig.4A for all four HCP datasets. We found that the similarity between the matched states generated by both approaches is more than 99%, with any value L of more than five, and the results were reproduced across four HCP datasets. The estimated states with conventional kmeans and iSparse kmeans (L=6) are shown in Fig.4B for all HCP datasets.

**Fig.4.**
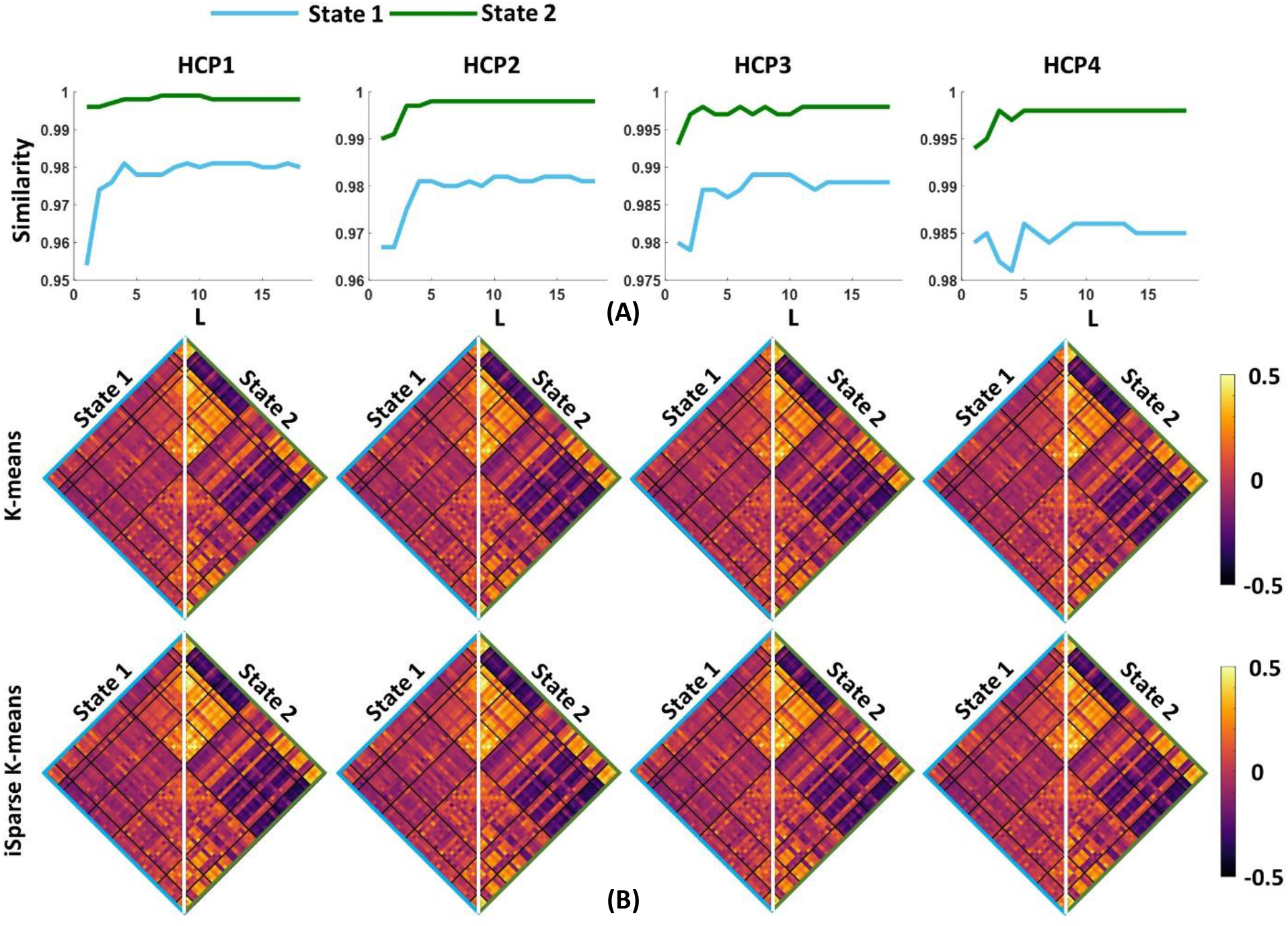
The estimated dFNC states with iSparse and conventional kmeans for all HCP datasets. (A) We swept the L value in the first kmeans clustering and calculated the similarity between the estimated states with iSparse and conventional kmeans. For any L>5, we did not find a significant improvement in the similarity between two clustering methods. (B) both iSparse and conventional kmeans generated similar dFNC states in all four HCP datasets.

### 3.2. iSparse kmeans finds the optimum cluster number faster than the conventional kmeans

After finding the minimum reliable value of L, we assessed the speed of our method in finding the optimum number of clusters and compared it with the conventional method when it uses the whole dataset. We evaluated the speed of our process with different percentages of data. The results are shown in Fig. 5A, B, C, and D for HCP1, HCP2, HCP3, and HCP4, respectively. We found that iSparse kmeans is faster when we use a lower percentage of data in each iteration, while the similarity between the matched states estimated with both standard kmeans and iSparse is still more than 98%. Additionally, our proposed method is 27 times faster in funding the cluster order than the traditional method when we use only 0.12% of data (one subject) in each iteration.

**Fig.5.**
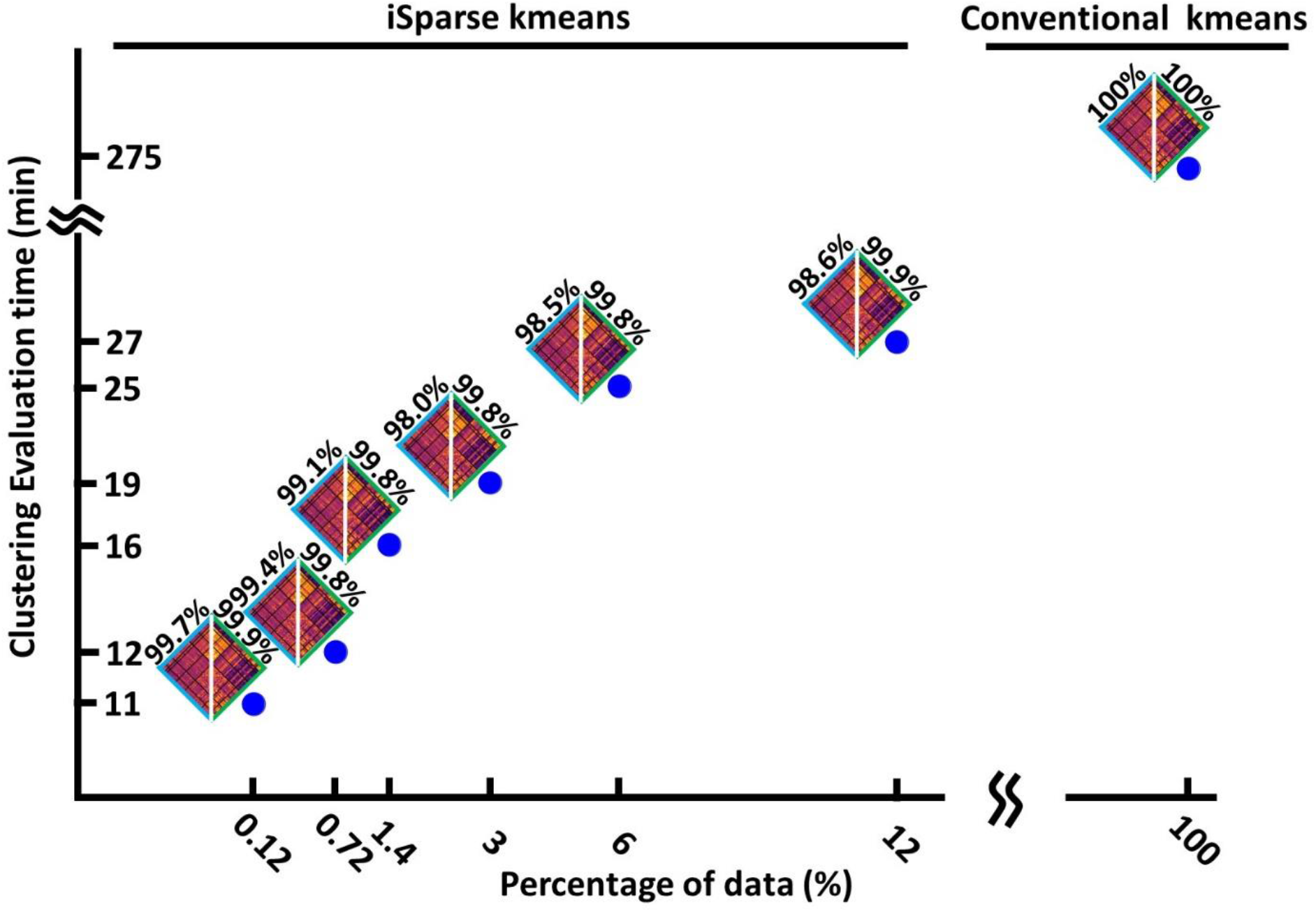
The clustering evaluation time with conventional and iSparse kmeans method. Reducing the percentage of the data used in each iteration of the first step, reduces the evaluation time. The iSparse kmeans method is 27 times faster the conventional method. The estimated states and their similarity with states estimated from whole data are shown for each percentage of data.

### 3.3. iSparse kmeans and conventional kmeans generate similar dFNC features

The next question is whether both clustering approaches generate similar dFNC features or not. To assess this, we estimated occupancy rate (OCR), the proportional amount of time each participant spends in a specific state, and the number of between-state transition numbers, for each participant in both standard and iSparse kmeans. Both features are estimated from the state vector, which shows the state of the brain in a given time (Fig.3 step4). Then, to assess the similarity between two methods in estimated dFNC features, we calculated the correlation between the result of the two methods. The results are shown in Fig. 6 A and Fig. 6B for OCR and the number of transitions, respectively, for all four HCP datasets. As Fig. 6A shows, the correlation between the estimated OCR by kmeans and iSparse kmeans is more than 0.98 (p<e^−10^). The result was replicated for all four HCP datasets. Additionally, the number of between-state transitions is significantly similar for both methods, and the result was repeated in all HCP datasets. This piece of evidence shows that our new clustering method produced similar dFNC features as well as the standard kmeans while our method is faster in finding the clustering order and does not require prohibitive levels of computational power.

**Fig.6.**
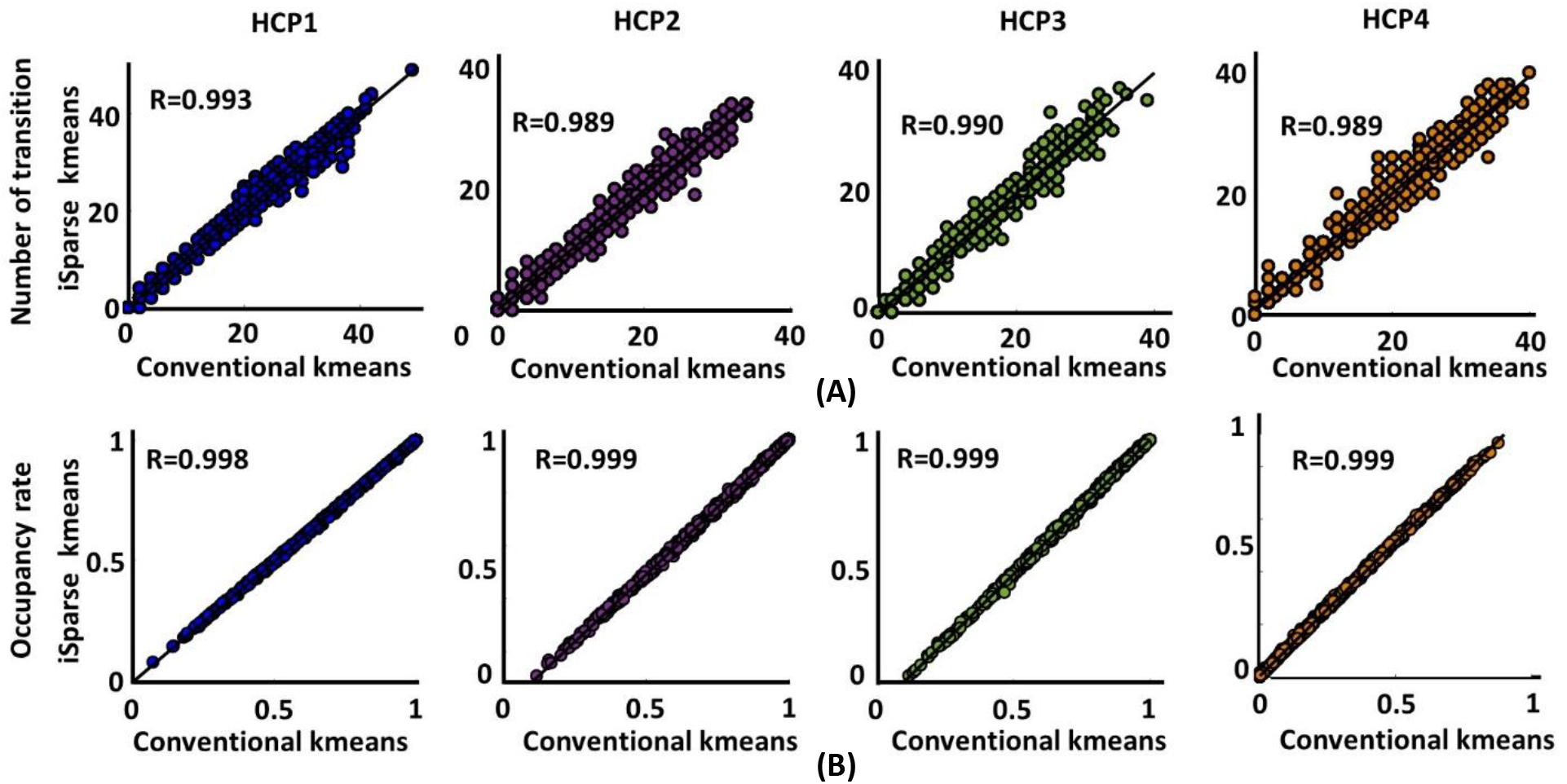
Both standard kmeans and iSparse kmeans generated similar dFNC features replicated across four datasets. (A) Estimated number of transitions from both standard kmeans and iSparse kmeans for all HCP datasets. The similarity between the estimated number of transitions from both method is more than 0.989. (B) Estimated occupancy rate (OCR) from both standard kmeans and iSparse kmeans for all HCP datasets. The similarity between the OCR from both method is more than 0.989 (*p*<0.0001, *N*=833).

### 3.4. iSparse kmeans has better cluster quality than the standard kmeans

Fig. 7 shows the distance ratio of both standard and iSparse kmeans for the optimum *k*=2 values in all four HCP sessions. We used a two-sample ttest to compare the distance ratio of standard kmeans vs. iSparse one. We found iSparse kmeans would have better cluster quality than the standard one by having a higher distance ratio (*p*<0.001, *N*=833).

**Fig.7.**
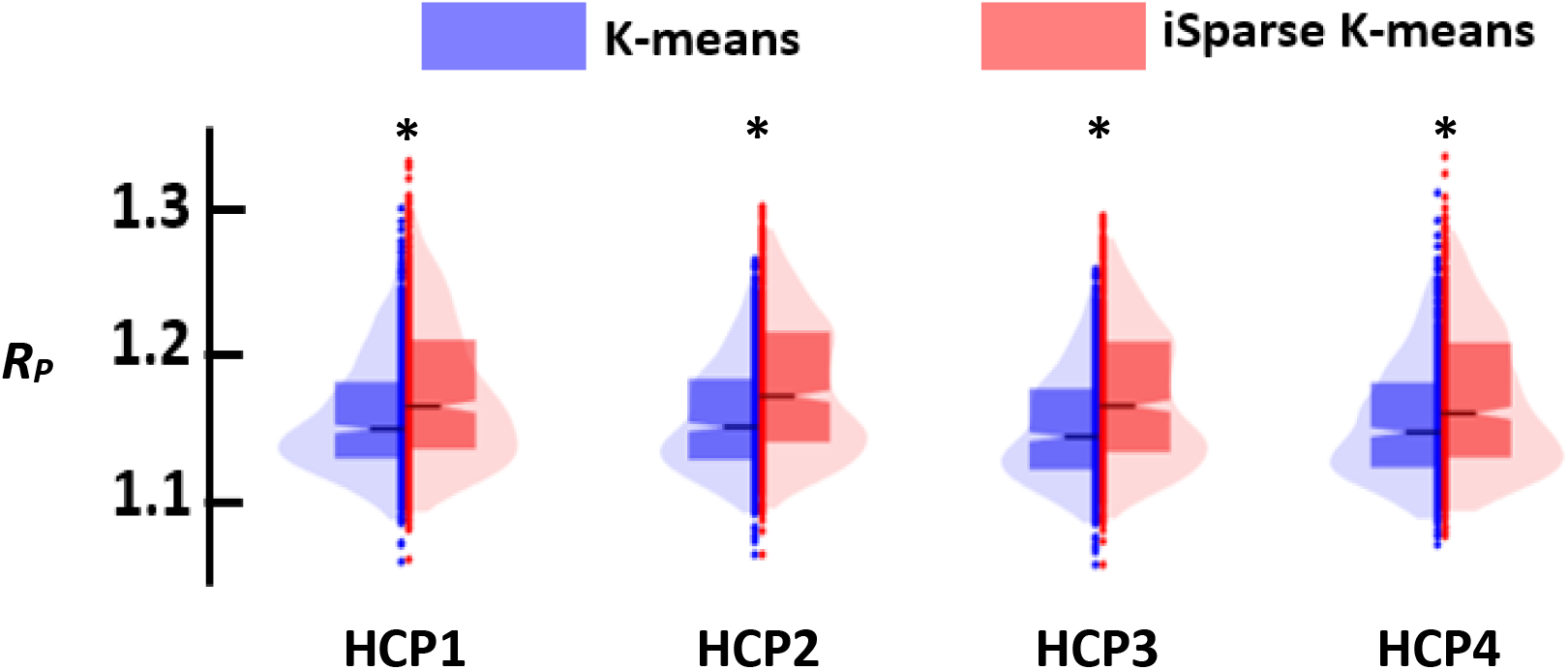
The comparison of the cluster quality between standard (blue) and iSparse kmeans (red) approach. Each columns represents the result for different k (cluster order). Each raw represents that result of each session. In all comparisons, iSparse kmeans had higher cluster quality (*p*<0.001, *N*=833).

## 4. Discussion

In this study, we developed an analytic pipeline to analyze large data dFNC information even without having a sophisticated computational resource. There are a few benefits of using this novel framework. 1) in the standard kmeans approach, we need to load the entire dataset, which can be computationally demanding and slow when using a large dFNC dataset. Our proposed method does not require loading the entire dataset. This dramatically reduces the required computational resources, 2) we showed our method is 27 times faster than the standard method in finding the cluster order, 3) we validated the reproducibility of the result across four sessions of rs-fMRI data within a population group; and 4) we demonstrated that our approach generates improved clustering quality compared to the standard approach.

Unlike standard kmeans, in which we need to load the entire dataset, our approach loads a portion of the data in each iteration. Therefore, we reduce both the required memory as well as the computational time. In this respect, our proposed algorithm is similar to mini batch kmeans, which partially loads the data and does not need expensive computational resources. But as (Béjar Alonso, 2013) shows, the cluster quality for mini batch kmeans is reduced compared to standard kmeans clustering, especially when the number of clusters increases. Unlike the mini batch kmeans approach, iSparse kmeans reduces the entire clustering process time (Fig.5) and increases the clustering quality (Fig.7).

Recent approaches for kmeans clustering of big data have focused on identifying the most informative features for the dataset and then running a kmeans on the reduced set. For example, a recent study reduced the dimension of the data set from *p* to *m* (*p*>*m*) by applying a principal component analysis on the entire dataset followed by a kmeans clustering on the projected dataset (Feldman et al., 2013). This method still needs the whole dataset to be loaded, which requires massive computational power. Additionally, since the kmeans is applied to the project space, we do not have an estimation of the cluster centroid in the original space. However, we can transfer the cluster centroid to the original space, but this estimate is inaccurate and yield lower cluster quality than the standard kmeans approach.

Our dFNC pipeline is based on the Neuromark pipeline, a fully automated independent component analysis (ICA) framework that uses spatially constrained ICA to estimate components that are flexible to each subject’s data and comparable across individuals (Du et al., 2020). Using the Neuromark pipeline, we calculated the replicated independent components for four HCP sessions. Additionally, we showed that 1) both standard and iSparse kmeans generated similar dFNC states in each session of HCP data, 2) the brain states were replicated across all four sessions using both standard and the proposed kmeans clustering approach. The reproducibility of the result across four sessions assessed the robustness of the proposed dFNC pipeline.

There are a few limitations to this study. First, our clustering method is not limited to kmeans clustering. We can adapt other fast clustering approaches to this pipeline and further improve the computational speed. Second, we did not compare our method’s computational speed and clustering quality with other existing fast clustering approaches. However, unlike these fast methods, we showed our approach generated better quality cluster than the standard kmeans clustering method. Future study is needed to compare the results across multiple clustering approaches. Third, we did not propose an algorithmic approach to set the maximum L value (Fig. 3). Finding the optimum L values is done empirically by running the method multiple times to evaluate replicability at different values of L.

## 5. Conclusion

Previous dFNC analytics pipelines use standard kmeans clustering, which is ill-suited for big dFNC data. Here, we developed a new method called iSparse kmeans clustering that reduced the evaluation time for finding the cluster order while we only loaded a portion of the dataset through several iterations. Therefore, in our new method, we do not need access to a strong computational power, as we need in the standard way for an extensive dataset. We validated that our method produces similar brain states and dFNC features as the standard method. Additionally, we evaluated the reproducibility of results across four HCP young adult datasets, which showed high robustness of the proposed method.

## 7. Author contributions

Mohammad. S. E. Sendi developed the method, analyzed the data, and wrote the manuscript. Robyn L Miller developed the method and provided feedback on the manuscript. Vince D Calhoun supervised the study and provided feedback on the manuscript.

## 8. Acknowledgments

We thank who participated in the HCP study and collected the data. This study was in part funded by NSF #2112455 and NIH R01MH123610.

## 9. Code and data Availability

The code and datasets used for this study are available on request to the corresponding author.

